# Model Annotation and Discovery with the Physiome Model Repository

**DOI:** 10.1101/498501

**Authors:** Dewan M. Sarwar, Reza Kalbasi, John H. Gennari, Brian E. Carlson, Maxwell L. Neal, Bernard de Bono, Koray Atalag, Peter J. Hunter, David P. Nickerson

## Abstract

**Motivation:** Mathematics and physics based computational models have the potential to help interpret and encapsulate biological phenomena in a computable and reproducible form. Similarly, comprehensive descriptions of such models encoded in computable form help to ensure that such models are accessible, discoverable, and reusable. To this end, researchers have developed tools and standards to encode mathematical models of biological systems enabling reproducibility and reuse; tools and guidelines to facilitate semantic description of mathematical models; and repositories in which to archive, share, and discover models. Biologists and clinicians can leverage these resources to investigate specific questions and hypotheses.

**Results:** We have comprehensively annotated a cohort of models with biological semantics. These annotated models are freely available in the Physiome Model Repository (PMR). To demonstrate the benefits of this approach, we have developed a web-based tool which enables users to discover models relevant to their work, with a particular focus on epithelial transport. In helping a user to discover relevant models this tool will provide users with suggestions for similar or alternative models they may wish to explore or utilize in their model based on the models they would like to use. The semantic annotation and the web tool we have developed is a new contribution enabling scientists to discover relevant models in the PMR as candidates for reuse in their own scientific endeavours. We believe that this approach demonstrates how semantic web technologies and methodologies can contribute to biomedical and clinical research.

**Availability and implementation:** https://github.com/dewancse/model-discovery-tool

## 1 Introduction

Over the years, computational models of human organ systems have been used to support and improve the diagnosis, treatment and prevention of diseases. These computational models help scientists to investigate specific research questions and hypotheses which may be ethically inappropriate, expensive to study experimentally, or not practically achievable using human or animal subjects. Discovery and exploration of relevant computational models can aid scientists to more quickly and accurately test their clinical or experimental hypotheses which may involve a broad range of biophysical mechanisms and observations such as disease states, drug actions and vast number of clinical observations. Ultimately this is expected to improve the quality, safety, effectiveness and reduce the spiraling cost of healthcare.

The IUPS Physiome Project (Hunter and Borg, 2003) and the Virtual Physiological Human (VPH) initiative (Hunter *et al*., 2010) aim at leveraging mathematics and physics based computational models to represent functional systems of the human body in a computable form, and most importantly, apply our gained physiological insight into clinical practice. In order to achieve this, reproducibility and reuse of computational models are key for building such computational models (Nielsen and Nickerson, 2017; Neal *et al*., 2014), as well as tools and open source software with the aid of modern technologies (Nickerson *et al*., 2016b; Garny *et al*., 2010; Cooper *et al*., 2010).

A key aspect of the Physiome Project and the VPH is to incorporate semantic annotation of the computational models such that these models are accessible, comprehendible, reusable, and discoverable. Here we specifically use semantic annotation to refer to the association of biological knowledge with the relevant constituents of a mathematical model in a computable and reproducible form. Such annotation does not, however,contribute to the mathematical content or interpretation of the model (Lister *et al*., 2009). The biological knowledge added in this way consists of a wide range of contextual information such as species, genes, physiological processes, and anatomical locations. Integration of such information in the computational models requires a diverse set of reference ontologies and often combines concepts from multiple ontologies, i.e. composite annotation (Gennari *et al*., 2011). Manual annotation is a time consuming process. We therefore utilize SemGen (Neal *et al*., 2018b), a semantic annotation tool, to annotate our cohort of exemplar models. To achieve this, SemGen combines concepts from a range of ontologies (Neal, 2010).

In order to discover relevant models, it is essential to ensure that models and their semantic annotations, along with other included metadata (e.g. authorship, original publication, etc), are findable and accessible by both people and computational services. This could then be used to identify missing resources and/or to add more information in the existing resources such that these are accessible, persistent, and discoverable. In this manuscript, we show model discovery within the Physiome Model Repository (PMR) (Yu *et al*., 2011), but more broadly, our approach could be applied to a variety of model repositories or resources.

Knowledge discovery is essential within the pharmacology discipline. A recent initiative, OpenPHACTS (J. Williams *et al*., 2012; Gray *et al*., 2014), has used semantic querying to enable chemists to discover and explore chemical entities and/or structures. Similarly, Piñero *et al*. (2017, 2015) have developed a discovery platform to explore human diseases and associated genes, as well as other variants using the wealth of semantic resources now available. Pharmacogenomics (Dumontier and Villanueva-Rosales, 2009) and biology (Goble *et al*., 2007) disciplines heavily depend on knowledge discovery with a view to accessing and integrating the web of science resources.

In this paper we present a web-based tool which will allow the user to discover models using the semantic knowledge in PMR. Specifically, the user would be able to discover models encoded in CellML (Cuellar *et al*., 2003), an XML-based language used to describe models in the PMR, including some useful biological information encapsulated within them: species, gene, anatomical location, related apical or basolateral membrane, etc. In section 2, we discuss some standards that enables researchers and scientists to encode mathematical models of biological systems. In addition, we describe a repository, the PMR, which consists of about 800 CellML models with varying levels of annotation. Detailed analysis of the biological information as well as the web tool by which we retrieve this information is discussed in section 3.

Here we focus on models of epithelial transport as a way of scoping the implementation of our web application. The tools being assembled here into a workflow of model discovery and reuse are, however, of utmost importance when considering the increasingly vast amount of experimental and clinical data being made available. For instance, when considering epithelial Na^+^ transport, retrieving multiple computational models of the epithelial Na^+^ channel (ENaC) would be important when considering new experimental data of normal and pathophysiological ENaC channel function since not all channel models may be able to represent the range of function observed. The ability to objectively select appropriate models both semantically and computationally and merge them across multiple scales of physiological complexity will allow researchers to build models that faithfully represent function and can reach up to upper level phenotypic measures observed in the clinic.

## 2 Tools and Standards

Computational tools and standards (Nickerson *et al*., 2016a,b) have evolved over the years to store, exchange and comprehend computational models, including their comprehensive descriptions, i.e. semantic annotation, such that knowledge resources can be accessible and discoverable. Some of the software, tools and standards used in our model discovery approach are presented below.

### 2.1 RDF and SPARQL

The Resource Description Framework (RDF) is a technology for encoding annotations (Klyne *et al*., 2018). RDF is machine readable and allows semantic interoperability between different systems. It represents annotations with a set of triples: subject, predicate, and object. The subject references the element being annotated, the predicate defines the assertion that is being made, and the object is the value being asserted. The COmputational Modeling in BIology NEtwork (COMBINE, 2018) community has recently recommended the use of RDF for storing annotations on models (Neal *et al*., 2018a).

SPARQL is a query language used to retrieve information from the RDF triples (Prud’hommeaux and Seaborne, 2018). SPARQL and RDF are both widely used semantic web technologies, which together enable “semantic querying”. Our tool relies extensively on the execution of semantic queries to discover knowledge.

### 2.2 CellML

CellML is an XML-based language that encodes mathematical models of biological phenomena (Cuellar *et al*., 2003). CellML models are modular and hierarchical, with model authors able to define abstract interfaces to their models by encapsulating complex details beneath a well defined interface. Model components can be reused with an import mechanism allowing external models to be accessible within the importing model. CellML includes MathML for representing equations; and makes use of the RDF for the description of metadata such as biological annotations. The CellML metadata specification (Cooling and Hunter, 2015) provides guidelines on how to annotate the biological semantics encapsulated in a CellML model using the RDF triples.

### 2.3 Ontologies

For our purpose, we have utilized the Chemical Entities of Biological Interest (ChEBI) (Hastings *et al*., 2015) to represent solute transports such as sodium, hydrogen; the Ontology of Physics for Biology (OPB) (Cook *et al*., 2011) to represent biophysical properties such as chemical concentration or fluid flow; and the Foundational Model of Anatomy (FMA) (Rosse and Mejino, 2003) to represent anatomical entities such as the cell nucleus and membrane. By utilizing the ChEBI, OPB and FMA, we can create composite annotations such as concentration of sodium in the collecting duct of the kidney. In addition, we have leveraged the European Bioinformatics Institute’s Ontology Lookup Service (OLS) (OLS, 2018) to elicit human readable labels from reference ontology URIs.

### 2.4 SemGen

SemGen is a toolset for annotating, merging, and decomposing models (Neal *et al*., 2018b). It is platform-independent and open source. It is capable of working with models encoded in a variety of modeling formats, including SBML and CellML. With SemGen, users can add composite annotations to a model that capture the biological meaning of a model’s contents (Gennari *et al*., 2011), including the variables in CellML models. These composite annotations link reference terms from knowledge resources such as the OPB, ChEBI, FMA, and GO to form precise, machine-readable descriptions of model variables.

The SemGen annotator interface of the NHE3 (Weinstein, 1995) model is shown in Fig. 1 where in the left panel, codewords and sub-models identify 39 CellML variables and 4 CellML components, respectively; and the right panel shows a composite annotation for flux of sodium from portion of renal filtrate in proximal convoluted tubule to portion of cytosol in epithelial cell of proximal tubule through sodium/hydrogen exchanger 3, as well as the declaration of that variable in the CellML model code.

**Fig. 1.**
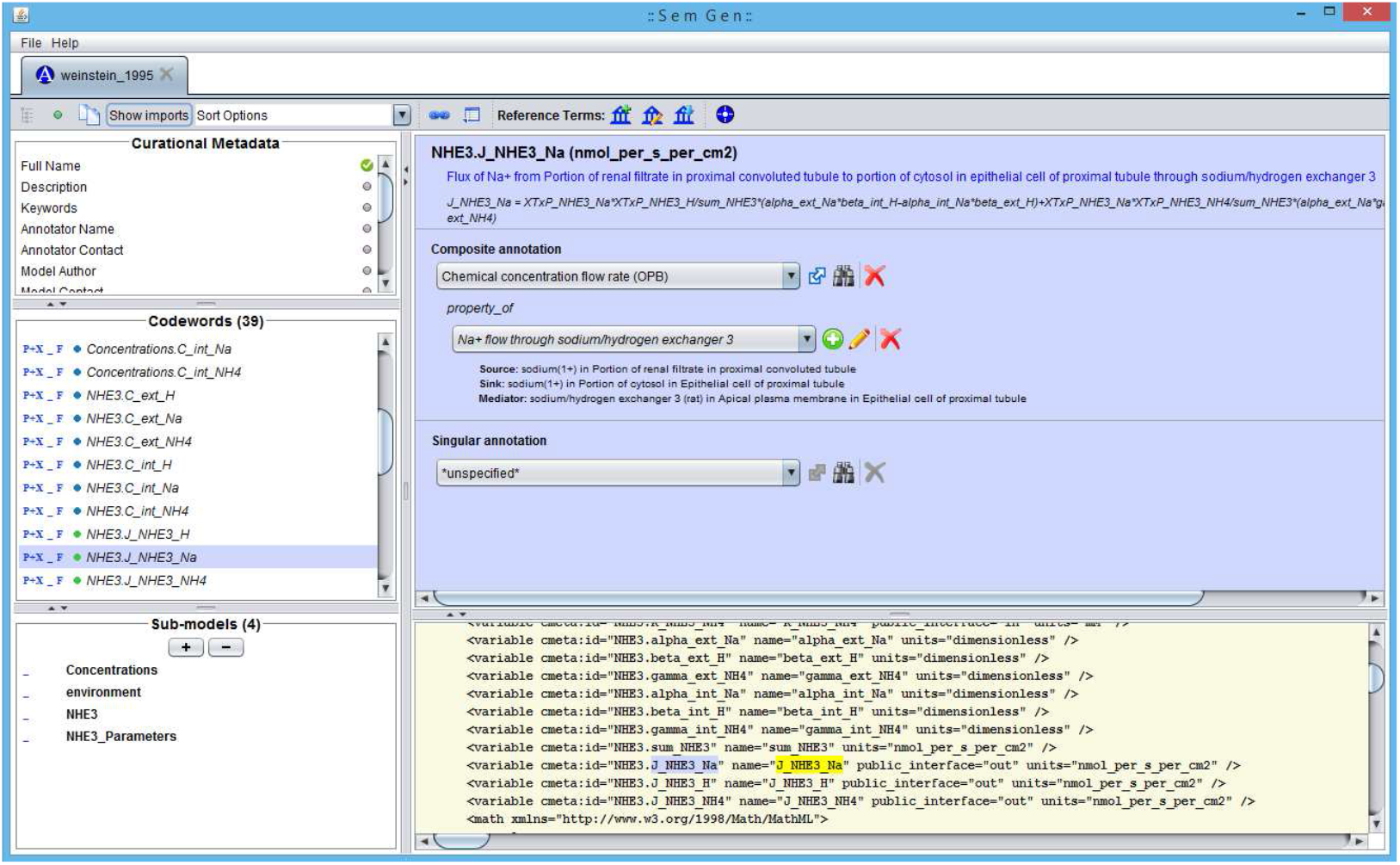
Semgen annotator interface of the Weinstein (1995) NHE3 model where codewords identify CellML variables and we have annotated the flux of sodium from portion of renal filtrate in proximal convoluted tubule to portion of cytosol in epithelial cell of proximal tubule through sodium/hydrogen exchanger 3.

### 2.5 The Physiome Model Repository

The Physiome Model Repository (PMR) was initially designed to store and manage CellML models (Yu *et al*., 2011) as part of the IUPS Physiome project (Hunter and Borg, 2003). In the PMR, CellML models can be annotated, exchanged, and reused. PMR supports a distributed version control system (DVCS) by which users can keep track of the changes made and thereby share and rollback to any committed point. By doing so, it enhances collaboration among model developers. Fig. 2 shows an example of synchronizing processes between client and server/cloud applications. In this case, tools like OpenCOR (Garny and Hunter, 2015) and SemGen maintain local copies of relevant files which are then pulled by the DVCS service (git) provided in the PMR. PMR has two primary data containers: workspaces and exposures.

**Fig. 2.**
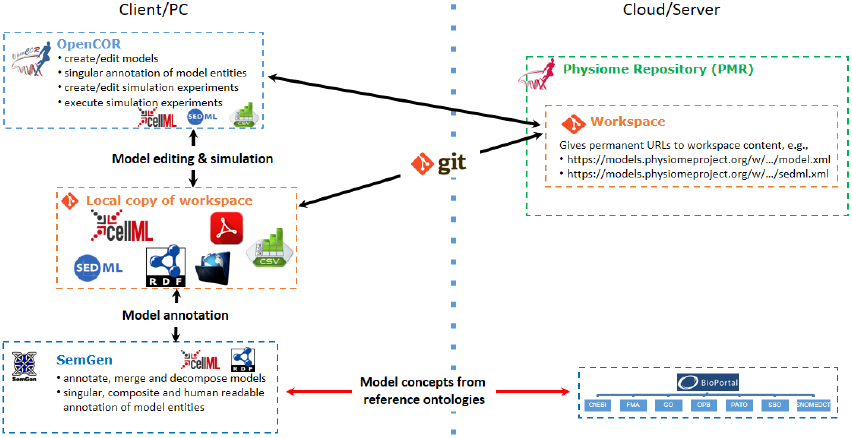
Schematic diagram of synchronization between PMR workspaces as well as OpenCOR and SemGen with the provision of git.

#### 2.5.1 Workspaces

A workspace is a DVCS repository for a user that contains related information in PMR. As discussed above, using a DVCS provides the ability to keep track of a number of commit points where users can easily switch to any specific point in the history of a workspace. A workspace can be shared with other users and thereby facilitates collaborative development. A workspace can be synchronized with other compatible DVCS repositories hosted anywhere on the Internet. Private workspaces can be made publicly accessible with the approval of a repository curator. A workspace can be assembled with another workspace, i.e. a hierarchical workspace, which resembles the modularity and reusability features of CellML. One of the key features in a PMR workspace is the indexing of RDF triples into an RDF store so that users can access these triples using SPARQL queries.

#### 2.5.2 Exposures

Exposures are equivalent to software releases – they provide a specialized view of a workspace at a specific point in the workspace’s history. Users can view the workspace in many ways dependent on the types of content in the workspace. For example, CellML models will have the following views generated: documentation; model curation and metadata; mathematics; source XML; and open with OpenCOR. Exposures are private by default and can be accessible after approval from a repository curator. Exposures can be migrated from one revision of the associated workspace to another as well as a separate “clone” of the workspace that another user might have created when reusing a model.

#### 2.5.3 PMR services

Web services address to the growing complexity of cross-platform applications though the exchange of messages encapsulated in the HTTP protocol. PMR uses web services and OAuth which maintains authentication of a user or consumer and manages access privileges for services provided by the PMR server. PMR web services exchange information as JSON formatted data and consumers have to follow specific guidelines (Yu, 2018) to fetch information from a PMR instance.

One of the services provided by PMR is a SPARQL endpoint, which can be utilised by application developers to submit SPARQL queries. Given the need for user access control and data persistence, PMR exposes a readonly SPARQL endpoint and any query attempting to modify the indexed RDF triples will be rejected.

## 3 Results

To demonstrate the value of the semantics-based approaches we propose here, we have developed a platform which allows the user to discover and explore models of interest and then display the annotated information via a web-based interface. We do this in the context of epithelial transport, although as mentioned above these technologies are broadly applicable across the spectrum of biomedical modelling. By restricting ourselves to epithelial transport we have been able to implement capabilities that we believe to be more useful in this specific domain. To provide an initial cohort of models and model entities for use in developing our tool, we first annotated the biological semantics of a set of models in the CellML format using the SemGen tool. These semantic annotations were deposited in PMR to provide resources to discover and assemble on the web interface. Detailed analysis of this process is described below and the demonstrator tool can be accessed via https://github.com/dewancse/model-discovery-tool.

### 3.1 Biological Annotation

An initial cohort of 33 models have been annotated with biological semantics and made available in PMR. This included 12 renal, 11 cardiac, 4 lung, 2 musculoskeletal and 4 miscellaneous epithelial transport models. In particular, we extracted each of the mathematical variables from the models and associated them with a biologically meaningful information so that computational modellers and clinicians can discover models relevant to their investigations. These models are all encoded in the CellML format and can be accessed via https://models.physiomeproject.org/workspace/527.

This initial cohort of well annotated models serves as a proof of concept for testing our implementation. According to the recent community agreement on the harmonization of annotation across the computational modelling in biology community (Neal *et al*., 2018a), we expect the repository of available annotated models to rapidly grow as the community begins to populate this repository. With community adoption of the harmonized annotation guidelines, we will be able to utilize our platform to query across all available repositories and model encoding formats.

Resources (variables, components, etc.) in these models can be uniquely identified with URIs composed of the CellML file name (including relative or absolute paths in the PMR) and the document-unique identifiers defined by the CellML format (so-called metadata identifiers). For example, when storing annotations in the same PMR workspace as the NHE3 (Weinstein, 1995) model, the URI `weinstein_1995.cellml#NHE3.J_NHE3_Na` unambiguously links to a variable defining the sodium flux through the NHE3 transporter. This example has been annotated in human readable text as “flux of sodium from portion of renal filtrate in proximal convoluted tubule to portion of cytosol in epithelial cell of proximal tubule through sodium/hydrogen exchanger 3”. Figure 1 shows the SemGen annotator interface for this example annotation formulated as a composite semantic annotation (Gennari *et al*., 2011). This demonstration combines concepts from multiple reference ontologies – OPB, FMA and the Protein Ontology (PR). The detailed composite annotation has been presented in Figure 3.

**Fig. 3.**
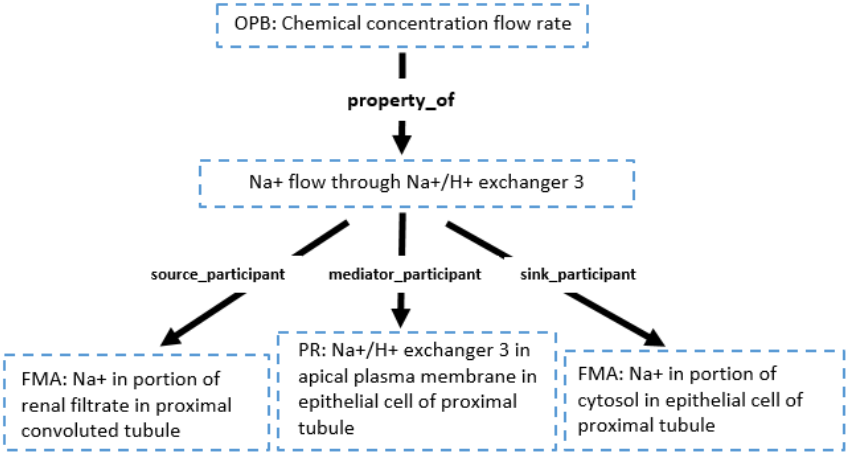
A composite annotation tree for flux of sodium from portion of renal filtrate in proximal convoluted tubule to portion of cytosol in epithelial cell of proximal tubule through sodium/hydrogen exchanger 3.

### 3.2 Web Based Tool

We have developed a web-based tool to discover and explore models of interest using information extracted from PMR. Figure 4 shows a list of discovered models for a search query “flux of sodium” from PMR. From this list, the user can investigate various options such as CellML model entity – name of the model, component name and variable name; biological meaning deposited in PMR; protein name; and species and genes used during experiments of the associated models.

**Fig. 4.**
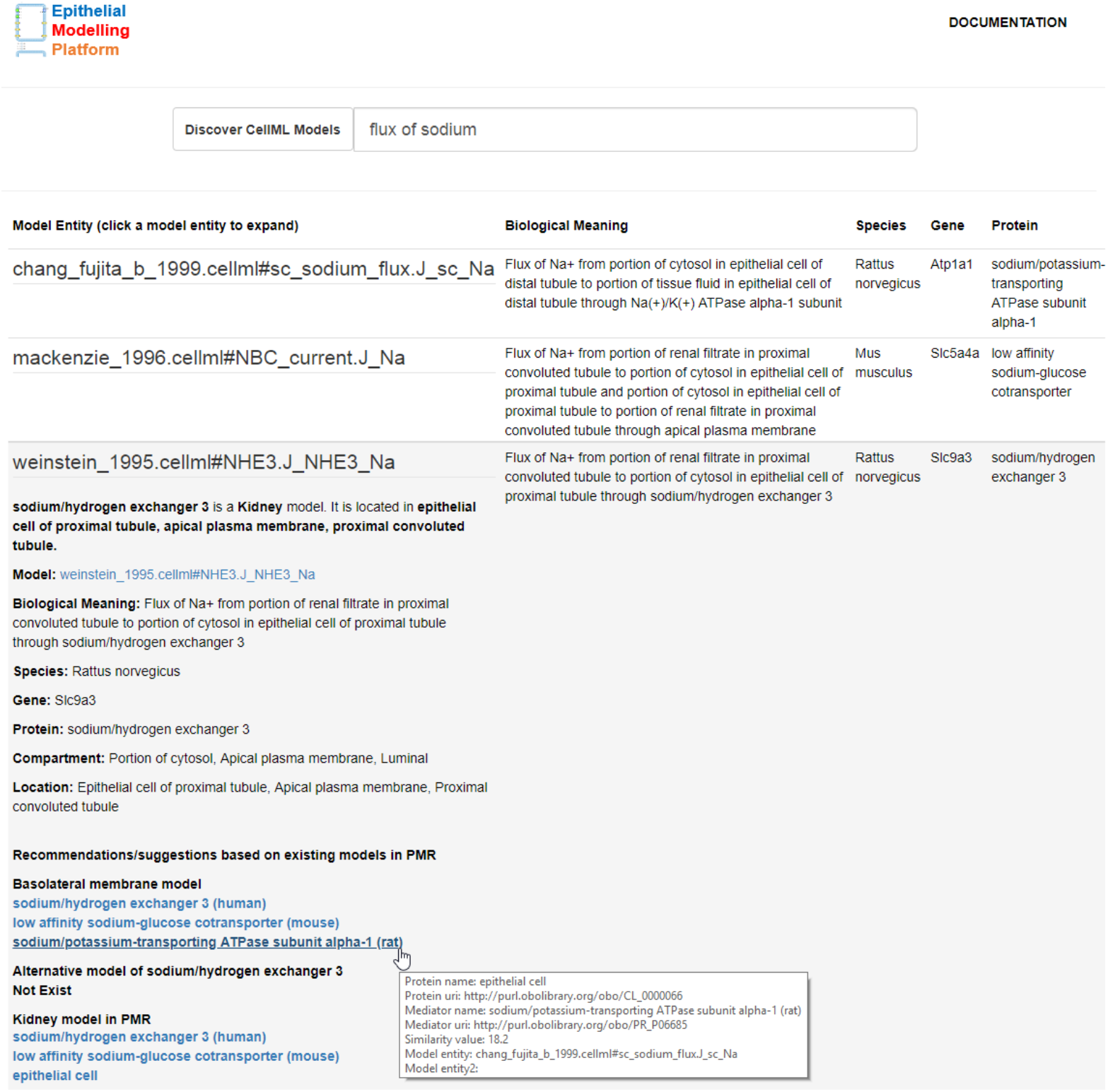
A use case application to search for models in PMR which are relevant to the query ‘flux of sodium’. By querying the knowledge PMR this tool retrieves components from the Weinstein (1995) NHE3 model, Mackenzie et al. (1996) SGLT2 model, and the Chang and Fujita (1999) epithelial cell model. Recommendations of models similar to the Weinstein (1995) NHE3 model are displayed as a result of the user selecting the NHE3 sodium flux in the returned results. See text for further details.

Users are able to enter human readable search terms and phrases or choose to enter specific vocabulary terms to search for. When discovering resources in PMR relevant to a given search phrase, we first decompose the English prose into a series of semantic queries which are executed on the PMR knowledgebase and ranked according to how well the discovered resource matches the initial search query. Continuing our example, the phrase shown in Figure 4 will best match any resource associated directly with a sodium flux, but other sodium-related resources will also be returned with a lower ranking in the search results. To achieve that, we have maintained a static dictionary with key and value pairs to map searched text to reference ontology terms. For example, for a search query “flux of sodium”, the tool will map flux and sodium to OPB and ChEBI ontology terms, respectively, as defined in the dictionary.

In addition, the user can click on a CellML model entity to explore more information as a recommendation or suggestion. As the name implies, this recommendation system provides the user with suggestions for similar or alternative models they may wish to explore or utilize in their model. Machine learning based recommender systems are widely used to build knowledge management systems enabling the user to interact with the system (Bobadilla *et al*., 2013; Almazro *et al*., 2010; Kywe *et al*., 2012; Carrer-Neto *et al*., 2012; Ricci *et al*., 1990; Wiesner and Pfeifer, 2014; Lémdani *et al*., 2011). However, our recommender system works in a different manner as it suggests relevant models’ description from the semantics in PMR, with respect to a model the user is interacting with on our platform.

Specifically, when the user clicks on a model, then the recommender system displays relevant models in PMR. Initially, it provides a brief overview of the selected model: name of organ and its anatomical location; biological meaning; species, gene and protein name; as shown in Figure 4. In this system, we have added a tooltip feature to quickly view some useful information such as protein name and its URI, mediator name and its URI if it is a co-transporter, CellML model entity, and a link to navigate to the associated CellML model.

This information is followed with relevant models in PMR with respect to the selected model. For example, if the selected model is located in the basolateral membrane, then existing basolateral model(s) will be populated as a suggestion for similar or alternative models. This populated list is ranked in an attempt to ensure more relevant models are closer to the top of the list.

As discussed above, we focus here on epithelial transport models which are often models of specific membrane-bound transport proteins. To determine which models in the repository are more relevant, we use the WSDbfetch (WSDbfetch, 2018) and Clustal Omega (Clustal, 2018; Sievers *et al*., 2011) services freely available from the European Bioinformatics Institute. For our purpose, we used WSDbfetch to retrieve a protein sequence for a given protein identifier, and the Clustal Omega service to obtain alignment metrics among multiple protein sequences calculated as a similarity matrix.

To get this matrix, we first fetched relevant protein models’ identifier located in the basolateral membrane. Next, we sent these protein identifiers to the WSDbfetch service to retrieve protein sequences. Finally, we sent these protein sequences to the Clustal Omega service to get the similarity matrix. An example of such a similarity matrix is shown in Fig 5. Along with the biological semantics, the similarity matrix provides a quantitative measure to help rank related models. In this manner, we hope to present the user with the most relevant models first. While these metrics are specific to membrane-bound transport proteins, we envision similar domain-relevant metrics can be developed as our methods are applied in the context of other domains.

**Fig. 5.**
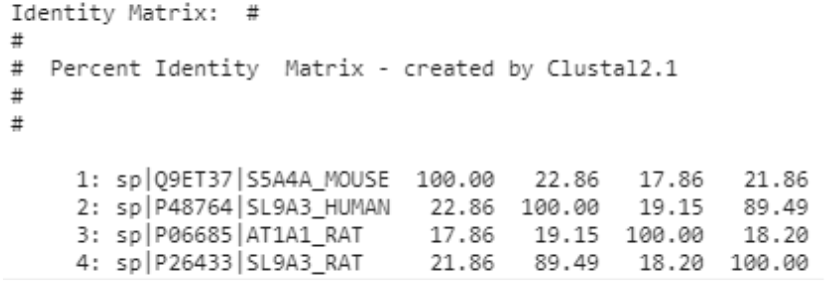
Similarity matrix from the EBI Clustal omega service where P26433 protein is the selected model and rest of the proteins are basolateral membrane models shown in Figure 4, see protein names under “Basolateral membrane model” section whose IDs and matrix scores are as follows: P48764 (89.49), Q9ET37 (21.86) and P06685 (18.20), with respect to the selected P26433 model.

An alternative option shown in Figure 4 is also provided for the user to explore the same selected model with different species and from various workspaces. The user could also explore relevant organ models and could navigate to the associated workspaces for further investigation.

## 4 Discussion

We have developed a novel semantic annotation and model discovery methodology with the aid of modern tools and standards and implemented a web-based tool for scientists to use in order to take advantage of this methodology. This semantic annotation uses composite annotation schema supported by the SemGen annotator tool. We have made available in PMR an initial cohort of models in the CellML format annotated with biological semantics to use in demonstrating the utility of our methodology and implemented platform.

In the model discovery approach, we have made cascading SPARQL calls to explore CellML models in PMR. First, it explores model name and biological meaning of a searched text. This text then maps to a static dictionary as key and value pairs with the help of reference ontologies such as OPB and ChEBI. Subsequent calls fetch the protein identifier of that model and we used this protein identifier to fetch the species and gene names of that model from the EBI ontology lookup service. A recommender system has been developed which provides useful information of a selected model from the web interface and suggests relevant models’ description from the semantics in PMR.

Alternatively, recent studies (Henkel *et al*., 2010; Köhn *et al*., 2009; Endler *et al*., 2009) have investigated machine learning techniques to extract and discover knowledge with a view to filtering search results. This enables similarities between models to be found as well as clustering models based on the semantic annotations. Combining information retrieval and clustering between models will enable scientists to efficiently explore and discover this knowledge (Schulz *et al*., 2011).

One of the limitations of the discovery approach is that we maintained a static dictionary with key and value pairs to map searched text to reference ontology terms, i.e. exact matching. However, this could be handled with Natural Language Processing (NLP) techniques described in (Sfakianaki *et al*., 2014, 2015) and by making SPARQL queries to the EBI SPARQL endpoint. For example, for a text “flux of sodium”, NLP will parse the text into pieces with tokenization, lemmatization, and part of speech techniques and then SPARQL queries to the EBI SPARQL endpoints will map these pieces into reference ontology terms. For convenience, the prototype application could also recommend similar words when the user misspells.

In addition to the discovery approach, we are moving forward to extend the web application into an SVG based Epithelial Modelling Platform for visualization and graphical editing based on biological semantics.

## Acknowledgements

We would like to thank the Semantics of Biological Processes lab at the University of Washington for valuable discussions and feedback during the April 2018 meeting in Seattle. The authors wish to acknowledge the Centre for eResearch at the University of Auckland for their help in facilitating this research. The availability of our live web application demonstration is made possible by use of the Nectar Research Cloud, a collaborative Australian research platform supported by the National Collaborative Research Infrastructure Strategy (NCRIS). Thanks to Sean Matheny at the Center for eResearch for his cooperation to setup our tool and service in Center for eResearch at the University of Auckland and Nectar.

